# A translational porcine model to assess the graded impact of hemorrhage and aortic occlusion on cardiovascular hemodynamics and renal perfusion

**DOI:** 10.64898/2026.02.05.703869

**Authors:** Fahim U. Mobin, Micaela K. Gomez, Sandra Januszko, Jacob Dooley, Antonio C. Renaldo, Heather Burkhart, James E. Jordan, Timothy K. Williams, Lucas P. Neff, Sadman Sadid, Matthew J. Eden, C. Alberto Figueroa, Elaheh Rahbar

## Abstract

Resuscitative Endovascular Balloon Occlusion of the Aorta (REBOA) is a lifesaving intervention used to manage non-compressible torso hemorrhage by temporarily occluding the aorta to minimize blood loss and preserve perfusion to vital organs. Partial REBOA (p-REBOA) has been proposed to mitigate ischemic injury associated with full-REBOA (f-REBOA). However, implementation of p-REBOA clinically has been challenging due to our limited understanding of the acute hemodynamics with p-REBOA particularly in relation to cardiac, carotid, and renal perfusion. In this study we developed and utilized a novel porcine model to continuously measure cardiac, carotid, renal and systemic hemodynamic responses to varying degrees of hemorrhagic shock and aortic occlusion. Yorkshire pigs (N=54) underwent instrumentation for continuous hemodynamic monitoring and hemorrhage was induced for 30 minutes to achieve 10%, 20%, or 30% blood volume loss (n=18/group), followed by randomized treatments of either no occlusion, p-REBOA, or f-REBOA occlusion strategies (n=6/group) for 30 minutes. After occlusion, shed blood was re-transfused over 15 minutes, and REBOA balloons were deflated and removed. This was followed by a 3-hour automated resuscitation and critical care period. Renal and carotid perfusion decreased progressively with hemorrhage severity. Interestingly, 30 minutes of f-REBOA resulted in significant ischemia-reperfusion injury where renal perfusion was profoundly suppressed to 40% of baseline renal flow. On the other hand, p-REBOA yielded superior renal perfusion, while maintaining cardiac function and carotid perfusion. p-REBOA also required less fluid and vasopressors. This translational pig model offers new opportunities to assess acute cardiovascular hemodynamics during interventions for the management of hemorrhagic shock.

## Introduction

Hemorrhagic shock remains the leading cause of death following a traumatic injury in individuals aged 1-44 years old.^3–6^ Non-compressible truncal hemorrhage (NCTH), in particular, contributes to the majority of these hemorrhage-related deaths, often within 1-6 hours post injury. This necessitates delivery of effective lifesaving interventions to manage NCTH and prevent associated hemorrhage deaths. Endovascular hemorrhage control strategies such as Resuscitative Endovascular Balloon Occlusion of the Aorta (REBOA) have increasingly been adopted for the clinical management of NCTH. During REBOA, a balloon catheter is introduced via the femoral artery and positioned within the thoracic or abdominal aortic region to temporarily occlude aortic flow, thereby providing short-term hemodynamic stabilization to vital organs (i.e., heart and brain) until definitive hemorrhage control can be achieved.^7^

Despite some successes and temporary benefits, implementation of REBOA has been associated with significant negative sequelae. Complete or full aortic occlusion with REBOA imposes an ischemic burden distal to the site of occlusion. This includes renal dysfunction, ischemia-reperfusion injury and lower extremity ischemia, especially when REBOA is deployed for prolonged durations^8,9^. These negative consequences have fueled ongoing debate among clinicians regarding the net clinical benefit of REBOA. For instance, some observational studies have reported increased patient mortality compared to non-REBOA patient populations with similar profiles^10^. As a function of this, the organ-specific physiological effects of REBOA, particularly on the heart, brain, and kidneys remain only partially understood.

Left ventricle pressure-volume (P-V) relationships are the gold standard in evaluating cardiac mechanics and heart function. P-V loops provide a cyclical relationship between key pressure and volume dynamics cardiac performance. Some common parameters extractable from P-V loops include cardiac output (CO), ejection fraction (EJF), arterial elastance (Ea), stroke volume (SV) and stroke work (SW), all of which work to that demonstrate changes in preload and afterload, reflecting the heart’s interactions with the systemic vascular conditions^11–14^. During hemorrhage, the cardiovascular system activates compensatory mechanisms to preserve organ perfusion. The heart responds to these needs by increasing heart rate, while systemic vasoconstriction works to modulate both vascular resistance and compliance. Given this dynamic physiological response, P-V data aid in assessing the hearts adaptation to progressive blood loss and REBOA^15^. In this context, P-V data can guide real-time clinical decisions in managing hemorrhagic shock and optimizing resuscitative efforts.

The clinical adoption of REBOA devices has been somewhat limited due to its underlying physiological mechanisms and impact not completely understood. Currently, most clinical studies on REBOA impact focus on its overall effect on hemodynamics and patient survival. While this approach yields a strong understanding of the global impact of REBOA, the dynamic nature of its impact on cardiac performance and organ-perfusion is comparatively less understood, a reality that becomes more complex considering the varying degrees of hemorrhage and their unique relationship with compensation. This leads into a lack of integrative physiological insight into the relationship between ventricular mechanics and their adaptation to the combined stresses of both hypovolemia and the afterload elevation induced by REBOA introduction, which impacts vascular perfusion, limiting the scope of occlusion strategy optimization to tailor the occlusion deployment across injury states.

Consequently, the complex interplay between hemorrhage severity, cardiovascular compensation, and resuscitation demonstrate a need for a more integrated physiological understanding. In this study, we exploited a novel translational porcine model with automated control to examine the impact of graded levels of hemorrhage and aortic occlusion. We hypothesized partial REBOA would be associated with superior hemodynamic support by preserving cardiac function, maintaining renal and carotid perfusion, and reducing ischemic burden compared to full REBOA strategies^16^.

## Methods

Yorkshire-cross pigs (N = 54; 27 females, 27 males; 6 per group) weighing 55–72 kg were obtained primarily from Palladium Biolabs, with limited sourcing from Oak Hill and Premiere BioSource early in the experiment. All procedures were approved by the Wake Forest University Institutional Animal Care and Use Committee (IACUC), and animals were acclimated for 72 hours prior to the start of experimentation.

Animals were fasted overnight and screened on the morning of each experiment. Exclusion criteria included white blood cell (WBC) count > 25 × 10^9^ cells/L or sustained pMAP < 20 mmHg. Anesthesia was induced via intramuscular Telazol® (5–7 mg/kg) and maintained with isoflurane (∼2%) following intubation. Mechanical ventilation-maintained end-tidal CO_2_ between a range of 35–45 mmHg, while normothermia (37–39°C) was preserved via an underbody warmer (BairHugger).

To counteract vasodilation induced by anesthetic isoflurane, norepinephrine (0.02–0.06 μg/kg/min) was intravenously administered to maintain MAP if 65 mmHg. Anticoagulation was supported by an initial heparin bolus (50 U/kg) through the left jugular vein at the end of instrumentation followed by continuous infusion (10 U/kg/hr) until the end of the experiment. 2 L of PlasmaLyte® was delivered during instrumentation, administered at ∼10 mL/kg/hr, and reduced to 5 mL/kg/hr prior to controlled hemorrhage. Basal NE infusion was maintained below 0.06 μg/kg/min prior to study initiation. Cumulative fluids and vasopressor use during instrumentation are reported in **Table 1**.

**Table 1.** Summary of maintenance fluid volumes and drug use during surgical instrumentation. Means and standard deviation are reported. Kruskal-Wallis test was performed to compare differences between hemorrhage groups. No statistically significant differences were observed, which means that our surgical instrumentation of all animals were consistent. Additionally, our baseline maintenance fluids, vasopressor and heparin use was also consistent across all animals.

|  | HEMORRHAGE LEVEL |  |  | Overall<br>(n=54) | p-value |
| --- | --- | --- | --- | --- | --- |
|  | 10%<br>(n=18) | 20%<br>(n=18) | 30%<br>(n=18) |  |  |
| **WEIGHT (kg)** | 62.6 ± 4.4 | 65.2 ± 4.9 | 64.4 ± 4.0 | 64.0 ± 4.49 | 0.19 |
| **AVG. ISOFLURANE (%)** | 1.8 ± 0.1 | 1.8 ± 0.1 | 1.8 ± 0.2 | 1.82 ± 0.15 | 0.2628 |
| **TOTAL INSTRUMENTATION TIME (minutes)** | 208 ± 27 | 215 ± 34 | 208 ± 32 | 210 ± 30.6 | 0.5696 |
| **AVERAGE RATE OF PLASMALYTE (mL/min)** | 20.9 ± 3.1 | 22.0 ± 2.7 | 22.1 ± 2.4 | 21.7 ± 2.75 | 0.3958 |
| **TOTAL PLASMALYTE VOLUME (mL)** | 4323 ± 622 | 4724 ± 829 | 4550 ± 611 | 4533 ± 701 | 0.3958 |
| **NOREPINEPHRINE/WEIGHT (mcg/kg)** | 5.71 ± 2.19 | 5.01 ± 2.75 | 5.40 ± 2.58 | 5.38 ± 2.49 | 0.67 |
| **CUMULATIVE NOREPINEPHRINE (mcg)** | 357 ± 135 | 332 ± 191 | 348 ± 169 | 346 ± 164 | 0.90 |
| **HEPARIN/WEIGHT (mL/kg)** | 0.092 ± 0.006 | 0.092 ± 0.006 | 0.092 ± 0.001 | 0.090 ± 0.008 | 0.09 |
| **CUMULATIVE HEPARIN (mL)** | 5.4 ± 0.5 | 6.0 ± 0.4 | 5.8 ± 0.6 | 5.82 ± 0.5 | 0.27 |

### Surgical Instrumentation

Surgical instrumentation of the animals was conducted in a similar fashion as previously published by our team and Sanin et al.^17^ Specifically, the left external jugular vein was surgically exposed and cannulated with a 7Fr triple lumen catheter for intravenous maintenance Plasmalyte (i.e., crystalloid infusion), medication infusion, bolus administration, and central venous pressure (CVP) monitoring via a fluid filled pressure transducer was obtained from the right jugular vein via a 7Fr sheath.

The left carotid artery was dissected, and a Transonic flow probe (4mm, Transonic, Ithaca, NY) was placed for carotid flow assessment. The left subclavian artery was surgically exposed and cannulated with a 7Fr sheath that extended into the descending thoracic aorta for proximal arterial pressure monitoring. Arterial sheaths were introduced into both the right (7Fr) and left (9/11Fr) femoral arteries. The aortic balloon catheter was inserted via the left femoral artery, and distal arterial blood pressure monitoring was via the right femoral artery. The left femoral vein was cannulated with a 9Fr multi-lumen access catheter for controlled venous hemorrhage, shed blood re-transfusion, and fluid boluses during the critical care resuscitation phase.

Once proximal pressure and venous lines were placed, a mid-line laparotomy incision was performed to remove the spleen. The splenectomy was performed to prevent autotransfusion during shock. A cystostomy tube was placed for urinary output measurement, and the left renal artery was dissected for flow probe placement. The right brachial artery was exposed and cannulated with a 5Fr micropuncture sheath for serial arterial blood sampling throughout the experiment. Finally, a sternotomy was performed to expose the heart for instrumentation of the flow probes at the aortic root and brachiocephalic arteries. A P-V catheter was introduced into the left ventricle via an apical stick.

### Flow Probe and Pressure Transducer Placement

To monitor the hemodynamic response, we used a Perivascular Flow Module with six ultrasonic flow probes (TS420, Transonic Systems Inc., Ithaca, NY). The six flow probes were placed at the aortic root (Ao, 22-24mm), brachiocephalic artery (BCA, 8mm), carotid artery (CA, 4mm), proximal descending aorta (PDA, 12mm), which was immediately proximal to the location of the balloon, renal artery (RA, 3mm), distal descending aorta (DDA, 6mm) before the femoral trifurcation, as illustrated in **Figure 1A**. Pressure transducers were placed in the left subclavian artery (LSA), DDA, and right jugular vein to obtain central venous pressure (CVP). Furthermore, an admittance pressure-volume catheter system (P-V catheter, ADV500 Transonic, Ithaca, NY) was inserted into the left ventricle via an apical stick for continuous pressure-volume (P-V) loop data capture, providing real-time left ventricular performance metrics. All hemodynamic parameters, including arterial pressures, volumetric flow rates, heart rate, cardiac output, and vascular resistance, were continuously recorded at 1,000 Hz.

**Figure 1.**
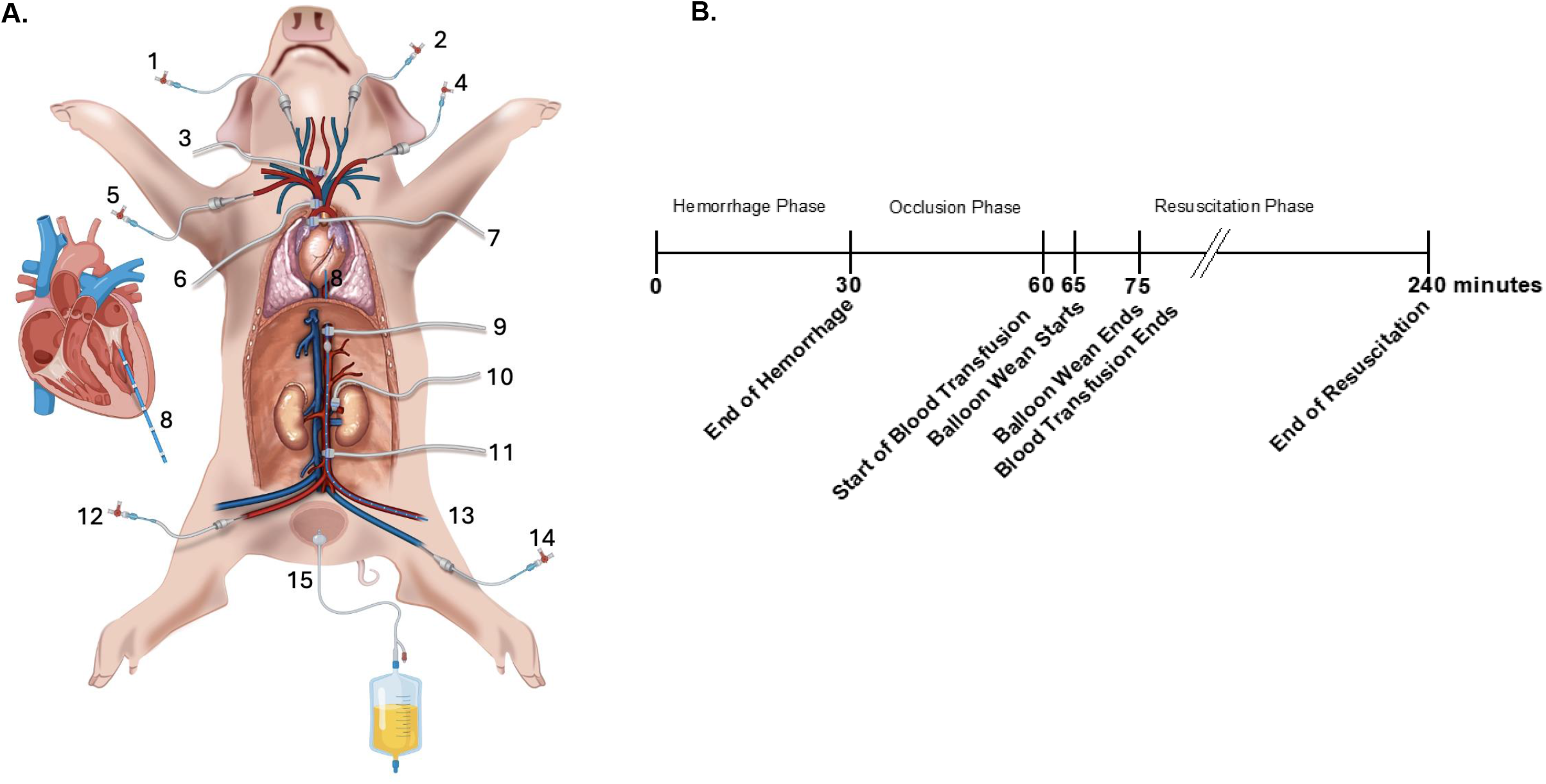
Illustration of animal instrumentation and experimental study timeline. **(A)** From superior to inferior and left to right: (1) Right External Jugular Vein – 7 Fr. cannula for central venous pressure (CVP) monitoring, (2) Left External Jugular vein – 7 Fr. triple lumen catheter for for fluid and drug administration, (3) Left Carotid Artery, 3-4mm Perivascular Transonic flow probe, (4) Left Subclavian Artery – 7Fr Cannula for proximal arterial pressure monitoring, (5) Right Brachial Artery, 5Fr. catheter for blood sampling, (6) Brachiocephalic arterial trunk, 6-8mm Perivascular Transonic flow probe, (7) Ascending Aorta (aortic root), 20-24mm Perivascular Transonic flow probe, (8) Left Ventricle, Transonic Pressure-Volume Catheter introduced via apical stick, (9) Proximal Descending Aorta, 12-14mm Perivascular Transonic Flow probe, (10) Left Renal Artery, 3-4 mm Perivascular Transonic Flow probe, (11) Distal Descending Aorta, 6-8 mm Perivascular Transonic flow probe, (12) Right Femoral Artery – 7 Fr. cannula reserved for distal pressure monitoring generally positioned infrarenal, (13) Left Femoral Artery - 9 Fr. cannula for introduction of the balloon catheter (balloon is inflated in the figure), (14) Left Femoral Vein – 9 Fr. MAC catheter for hemorrhage, transfusion and fluid resuscitation via automated Masterflex pump, and (15) Foley catheter for urine collection. Not depicted in the illustration are the endotracheal tube, splenectomy, or rectal temperature probe. We’d like to acknowledge Mr. Jerry Louis Shelton at Texas A&M University for his assistance in generating the pig illustration in panel 1A. **(B)** Experimental timeline depicting controlled 30-minute hemorrhage period, followed by 30 minutes of aortic occlusion, a 10-minute balloon wean and 3-hour critical care resuscitation phase. Animals were randomized to 10%, 20% or 30% hemorrhage (by volume), and then randomized to either no aortic occlusion, partial REBOA (p-REBOA) and full REBOA (f-REBOA) intervention groups. Autologous blood re-transfusion and balloon weaning occurred between 65–75 minutes, after which a closed-loop critical care algorithm was initiated to maintain target MAP>65 mmHg for three hours (until time 240 minutes).

As observed in prior studies, stroke volume dynamically varies during hemorrhage and resuscitation, requiring periodical admittance system recalibration^15,18^. A custom macro on LabChart was incorporated to computed cardiac output from integrated aortic root flow and derived stroke volume using heart rate extracted from arterial pressure waveforms. Manual device recalibration was performed every 5 minutes during hemorrhage and occlusion phases as well as at the start of each experimental phase thereafter.

Prior to the beginning of the study (T0) Animals were randomly assigned to experimental groups using a block randomization scheme. This method worked to ensure that an equal distribution of sex and hemorrhage severity was developed across treatment arms. Each subject experienced a unique hemorrhage severity over a 30-minute period: 10% low hemorrhage, 20% mild hemorrhage or 30% severe hemorrhage with total blood volume estimated relative to the assumption of 60 mL/kg.

After the hemorrhage phase ended (T30), the subjects were treated with various REBOA treatments, either no occlusion, partial occlusion, or full occlusion. This occlusion phase lasted for 30 minutes until T60. At T60, autologous blood re-transfusion began for a 15-minute period and 5 minutes after the start of this phase, a 10-minute gradual balloon weaning procedure was implemented; both of these ended at T75. Post-intervention care was managed using an automated critical care algorithm targeting MAP ≥65 mmHg. The experimental timeline is shown in **Figure 1B**.

p-REBOA was achieved in this study by algorithmic inflation of the balloon catheter to maintain a target distal-to-proximal pressure gradient while preserving measurable distal aortic flow. Balloon volume was titrated to maintain proximal MAP ≥ 65 mmHg while allowing continuous flow in the distal descending aorta as confirmed by flow measurements. f-REBOA conversely was defined as complete cessation of distal aortic flow. Post-intervention resuscitation after T85 was administered by an in-house proprietary critical care algorithm designed to maintain MAP ≥ 65 mmHg. The algorithm adjusted plasmalyte, norepinephrine and vasopressor infusion rates based on real-time measurements.

Left ventricular P–V loops were analyzed using a custom Python-based algorithm^15^. High-resolution data was processed using this algorithm to identify individual cardiac cycles, derive load-dependent parameters, and generate ensemble-averaged representative loops for each experimental phase. Hemodynamic and perfusion data were analyzed using ensemble-averaged measurements. Group differences were assessed using appropriate parametric or nonparametric tests, with two-way ANOVA models evaluating REBOA effects over time while adjusting for relevant covariates. Statistical significance was set at p < 0.05.

## Results

To ensure that surgical instrumentation time and perioperative fluids were consistent between all animals before the start of the experiment, we analyzed each variable from **Table 1** and observed no statistically significant differences between hemorrhage groups. The average instrumentation time was 210 ± 30.6 minutes and average rate of plasmalyte infusion was 21.7 ± 2.75 mL/minute. With regards to vasopressor usage, the average cumulative norepinephrine dose during instrumentation was 346 ± 164 mcg. A low dose of heparin was infused, totaling to 0.0898 ± 0.00815 mL/kg over the entire instrumentation time.

At the end of the hemorrhage phase (T30), increasing blood loss was generally associated with deterioration in cardiac performance as assessed by variations in the pressure–volume loop parameters and decreases in both renal and carotid flow as noted in **Table 2** and demonstrated in **Figure 2**. Cardiac output (CO) and end-systolic pressure (ESP) declined significantly with increasing hemorrhage severity. Notably, the most pronounced reduction in CO between hemorrhage levels was observed in the differential from 20% to 30% blood loss. Stroke work (SW) and stroke volume (SV) demonstrated statistically significant reductions observed at 30% hemorrhage compared to both 10% and 20%, while the differences between 10% and 20% hemorrhage were not significant. End-systolic volume (ESV) decreased significantly at 30% hemorrhage when compared to both 10% and 20% hemorrhage groups. This parameter shift implies impaired ventricular emptying at the high levels of blood loss. Ejection fraction (EF) was preserved at lower hemorrhage levels and demonstrated significant differences only between 20% and 30% hemorrhage. Heart rate (HR) increased from 10% to both 20% and 30% hemorrhages but remained stable between 20% and 30% hemorrhage, implying a stabilization in heart rate after mild hemorrhage. Arterial elastance (Ea) increased in hemorrhage states, while end-diastolic pressure (EDP), on the other hand, decreased significantly. Collectively, these findings demonstrate a graded decline in cardiac performance with increasing hemorrhage, characterized by preserved compensation at mild-to-moderate blood loss and marked decompensation at 30% hemorrhage, implying decompensatory effects being more apparent at higher levels of hemorrhage. Renal and carotid perfusion was also analyzed, and both exhibited nearly identical responses to increasing hemorrhage severity, reflecting a systemic reduction in perfusion as a function of changes in blood volume, consistent with increasing hemorrhage severity. This means renal and carotid flow decreased progressively as blood loss increased, as was expected. For these regions, all pairwise comparisons between average flow measurements across hemorrhage levels were statistically significant, indicating significant perfusion loss stratification with increases in hemorrhage severity.

**Table 2.** Baseline Cardiac and Hemodynamic Measurements. Summary statistics, reported as mean ± standard deviation, of cardiac output, heart rate, left ventricular (LV) pressure, proximal MAP and regional flow rates measured at the beginning of the experiment are shown. Using a Kruskal-Wallis test, no significant differences were observed across hemorrhage levels (10%, 20%, 30%) at baseline, indicating that all animals started with comparable cardiac and hemodynamic levels.

|  | HEMORRHAGE LEVEL |  |  | Overall<br>(n=54) | p-value |
| --- | --- | --- | --- | --- | --- |
|  | 10%<br>(n=18) | 20%<br>(n=18) | 30%<br>(n=18) |  |  |
| **BASELINE HEMODYNAMICS** |  |  |  |  |  |
| **CARDIAC OUTPUT (CO, L/min)** | 5.2 ± 2.1 | 4.3 ± 1.3 | 5.6 ± 2.0 | 5.0 ± 1.9 | 0.128 |
| **HEART RATE (HR, bpm)** | 97 ± 22 | 94 ± 18 | 105 ± 25 | 99 ± 22 | 0.444 |
| **END SYSTOLIC LV PRESSURE (mmHg)** | 71 ± 14 | 76 ± 7 | 73 ± 7 | 73 ± 10 | 0.186 |
| **END DIASTOLIC LV PRESSURE (mmHg)** | 4 ± 2 | 4 ± 1 | 4 ± 2 | 4 ± 2 | 0.927 |
| **PROXIMAL MAP (mmHg)** | 57 ± 5 | 61 ± 5 | 58 ± 7 | 59 ± 6 | 0.2 |
| **CAROTID FLOW (mL/min)** | 286 ± 85 | 309 ± 58 | 277 ± 88 | 291 ± 78 | 0.448 |
| **RENAL FLOW (mL/min)** | 215 ± 49 | 253 ± 60 | 230 ± 57 | 233 ± 57 | 0.186 |

**Figure 2.**
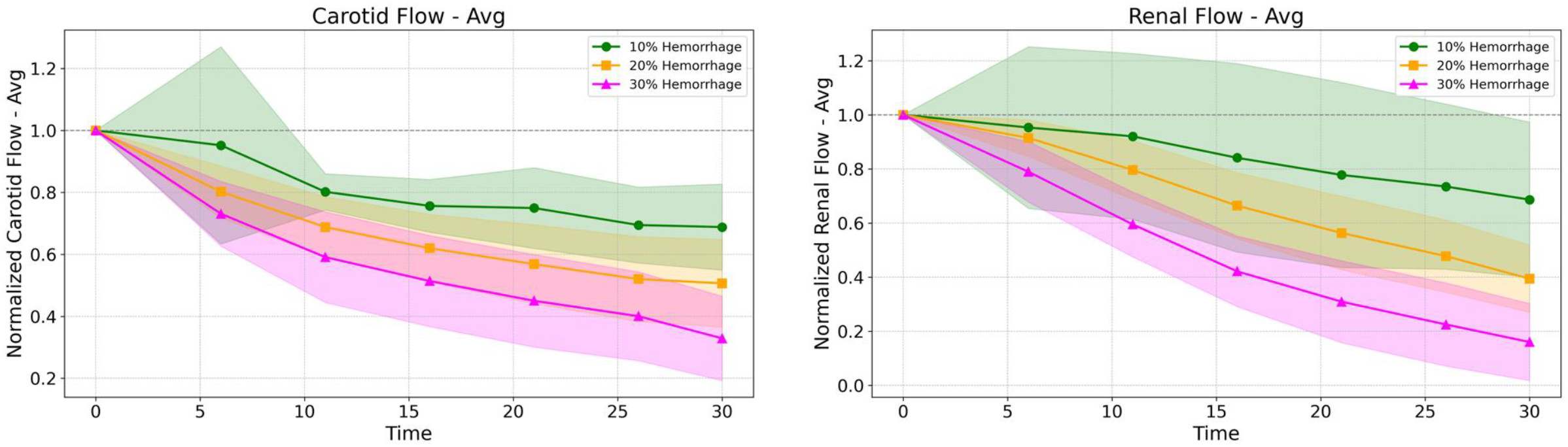
Normalized carotid and renal perfusion decline with increasing hemorrhage. Carotid and renal flow rates were normalized to each animal’s baseline value. Within each panel, mean data (N=18/group) is shown for 10% hemorrhage (green circle), 20% hemorrhage (yellow square) and 30% hemorrhage (pink triangle) groups over the 30-minute hemorrhage period. Shaded regions correspond to the 95% confidence interval. We observed a significant reduction in carotid flow (∼40%) was observed in animals undergoing 30% hemorrhage. Renal flow also declined with increasing hemorrhage, where animals experiencing 30% hemorrhage presented with 10-20% of baseline renal flow.

At the end of the occlusion phase (T60), both hemorrhage severity and REBOA occlusion type had significant impact on cardiac mechanics, as illustrated in **Figure 4**. For instance, Ea was high across all full occlusion subjects, but did not increase with hemorrhage severity, and was generally variable. Conversely ESV and EDV declined while when both stressors acted on the system, ESP increased and EDP decreased, indicating altered ventricular loading conditions. SW and SV declined significantly in cases of major 30% hemorrhage while lower 10% and moderate 20% hemorrhages did not express major differences, consistent with compensation effects, which led to the preservation of these variables. HR increased with hemorrhage but plateaued at higher levels of hemorrhage, while occlusion was associated with a reduction in HR. EF declined under severe hemorrhage levels and in both full and partial REBOA treatment groups. Overall, these data indicate that hemorrhage and occlusion exert substantial and, in in the case of some parameters, additive effects on ventricular performance. In the context of this study, this reveals the limits of cardiac compensation under severe physiological stress induced by a post hemorrhage state and occlusive REBOA treatment.

At T60, renal and carotid perfusion, illustrated in **Figure 3**, demonstrated divergent responses to hemorrhage and occlusion, in contrast to the trends observed with the cardiac performance parameters. Significant interaction effects between occlusion and hemorrhage were significant in that there was a positive impact on carotid flow, whereas renal blood perfusion was suppressed primarily by the presence of occlusion rather than by varying hemorrhage severity.

**Figure 3.**
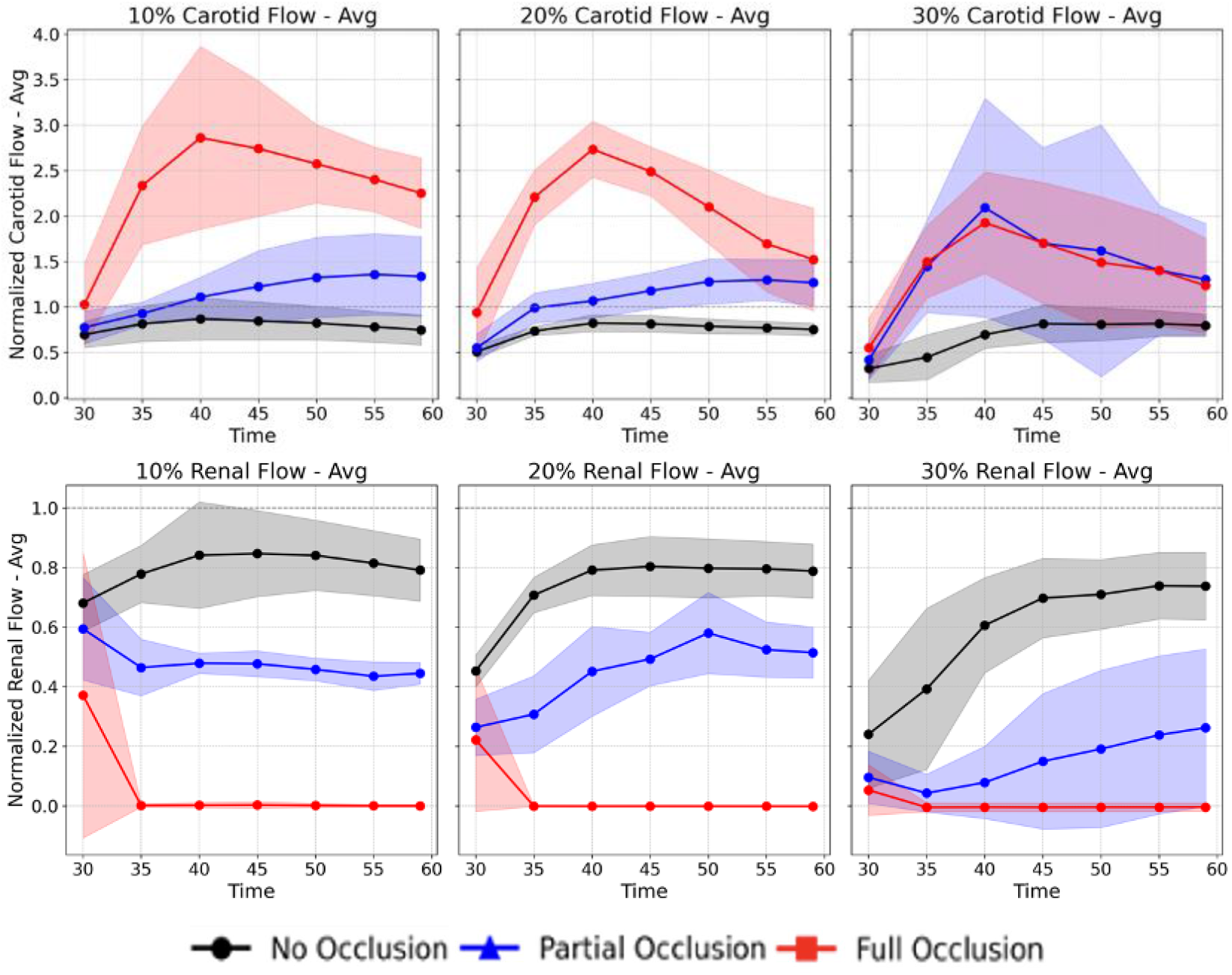
Normalized carotid and renal perfusion during degrees of aortic occlusion. Carotid and renal perfusion responses during the occlusion phase from T30 to T60, sampled at 5-minute intervals are shown. Flow rates were normalized to each pig’s baseline value. Significant interaction effects between degree of aortic occlusion and hemorrhage severity were observed, with carotid flow increasing up to 3x during full REBOA (f-REBOA), consistent with redistribution of blood toward the cerebral circulation. In contrast, renal perfusion declined during both partial REBOA (p-REBOA) and completed ceded in response to f-REBOA. Within each panel, perfusion responses are shown for no occlusion (black), p-REBOA (blue), and f-REBOA (red). Shaded regions correspond to the 95% confidence interval.

**Figure 4.**
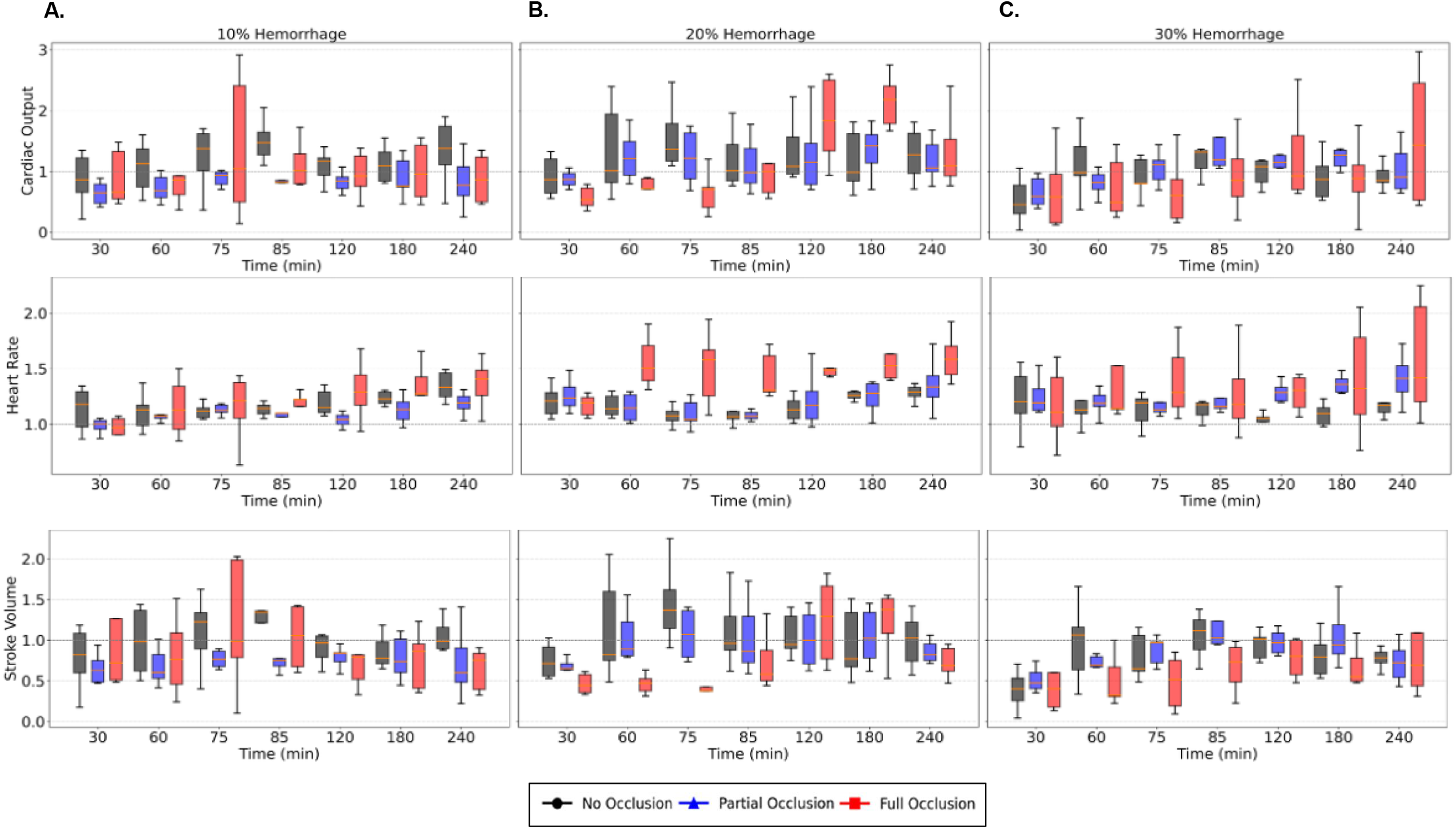
Cardiac response to graded levels of hemorrhage and aortic occlusions. Box plots illustrate normalized cardiac output (CO), heart rate (HR), and stroke volume (SV) measured across the entire experiment. All data are normalized to the subject’s baseline value. Data is presented at critical time points: end of hemorrhage (30 min), end of aortic occlusion (60 min), end of balloon wean (75 min) and during critical care resuscitation period (60-240 minutes). The first column (A) represents 10% hemorrhage, followed by (B) 20% hemorrhage, and (C) 30% hemorrhage. Within each panel, the aortic occlusion groups are color coded: no occlusion (black), partial REBOA (p-REBOA, blue), and full REBOA (f-REBOA, red). Increasing hemorrhage severity was associated with a graded decline in CO and SV, with the most pronounced reductions observed at 30% hemorrhage. As expected, HR increased with increasing hemorrhage and there was sustained tachycardia within the f-REBOA group at 20% and 30% hemorrhage.

Full occlusion induced a severe and consistent reduction in renal perfusion across hemorrhage levels, as was expected due to the mechanics of REBOA and its impact on the system. Carotid perfusion was impacted differently, increasing under occlusion, consistent with redistribution of blood toward the cerebral circulation, inherently demonstrating the intended goal of REBOA employ was met. Despite this, however, the benefit of increased perfusion diminished as hemorrhage severity increased. This indicates that hemorrhage-occlusion interactions were significantly impactful on carotid flow, indicating that the magnitude of occlusion-related augmentation depended on the levels of circulating volume after shock. These findings highlight a trade-off in perfusion during REBOA, demonstrated by either promotion or, at minimum, preservation of baseline cerebral blood flow at the expense of inducing renal hypoperfusion.

Fluid and vasopressor demands across the entire experiment were summarized and shown in **Figure 5**. Cumulative resuscitation volumes with Plasmalyte® were significantly higher in pigs that were randomized to full REBOA (f-REBOA), particularly post-20% and 30% hemorrhage. Vasopressin use was significantly greater in pigs randomized to p-REBOA and f-REBOA, regardless of hemorrhage severity. Norepinephrine requirements were highest in pigs who experienced 20% hemorrhage subjects and randomized to f-REBOA.

**Figure 5.**
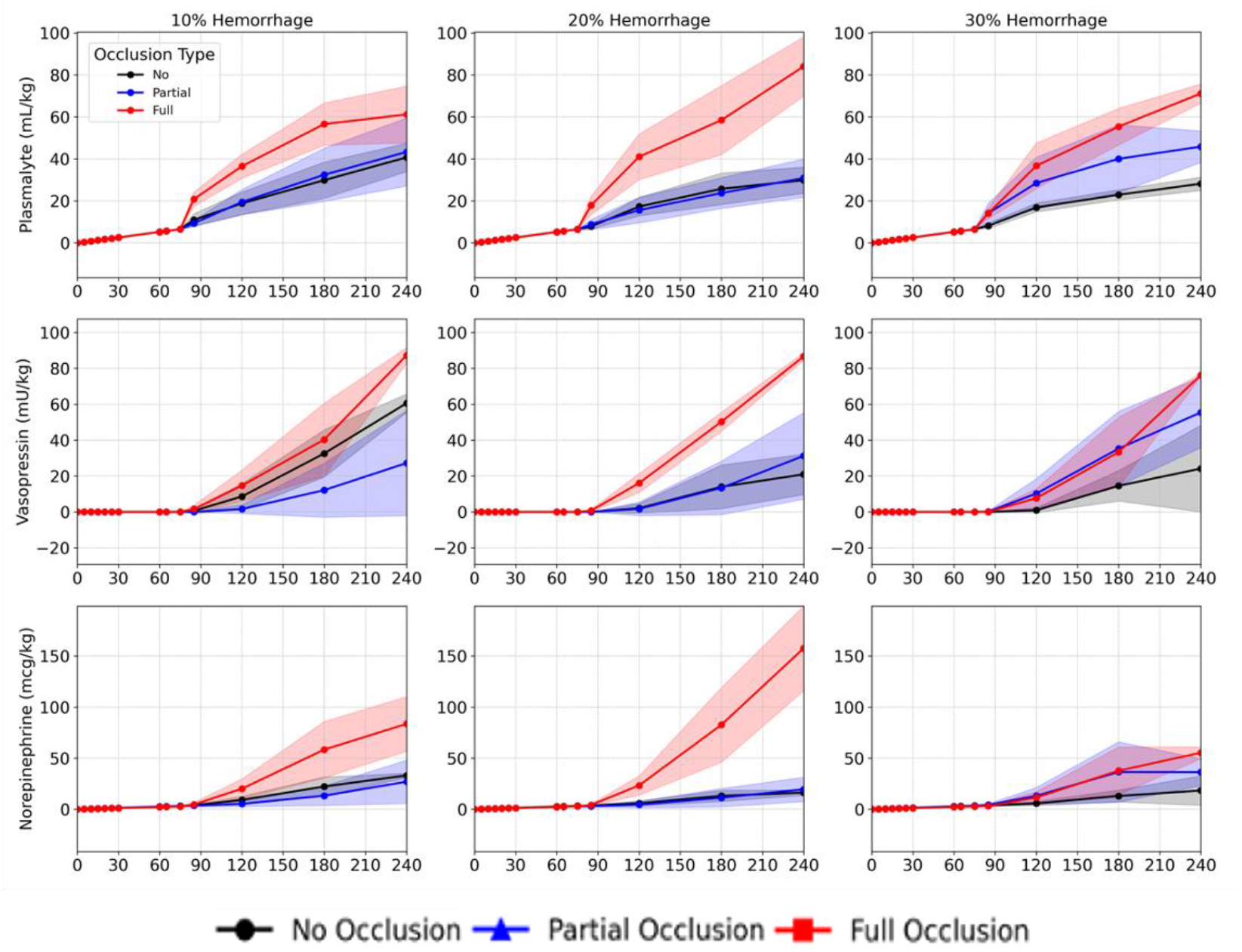
Fluid and vasopressor usage across hemorrhage severity and degrees of aortic occlusion. Fluid and vasopressor demands across the entire experiment are summarized. Cumulative resuscitation volumes with Plasmalyte® were significantly higher in pigs that were randomized to full REBOA (f-REBOA), particularly post-20% and 30% hemorrhage. Vasopressin use was significantly greater in pigs randomized to p-REBOA and f-REBOA, regardless of hemorrhage severity. Norepinephrine requirements were highest in pigs who experienced 20% hemorrhage subjects and randomized to f-REBOA. Within each panel, average cumulative volumes are shown for no occlusion (black), p-REBOA (blue), and f-REBOA (red) groups. Shaded regions correspond to the 95% confidence interval.

## Discussion

This study provides comprehensive and physiological characterization of controlled hemorrhagic shock independent of resuscitation as well as post-hemostatic REBOA intervention by combining high-fidelity pressure-volume analysis with regional carotid and renal perfusion over a variety of hemorrhage severities and occlusion strategies. Our study findings demonstrate that the cardiovascular system mechanistically exhibits robust compensatory behavior during hemorrhage with early signs of strain in moderate hemorrhage cases, but progresses toward marked decompensation at more severe hemorrhage levels, congruent what would be expected in cases of graded hemorrhage^1,2,15,18^. Importantly, while aortic occlusion imposes an additional physiological burden that alters cardiac mechanics, it redistributes blood flow in an organ-specific manner, revealing both the benefits and costs of occlusive resuscitation, an effect also observed in similar studies^15,16,19,20^.

At the end of hemorrhage (T30), all P-V parameter dynamics, with the exception of HR and Ea, demonstrated significant declines in cardiac performance as the severity of blood loss increased, with significant variations presenting at 30% hemorrhage. CO and ESP declined progressively with increase in hemorrhage, while SV, SW, EJF and ESV exhibited significant declines, especially at the highest hemorrhage level congruent with what was observed in a previous study^15^. This pattern suggests that compensatory mechanisms effectively preserve systolic function and ventricular emptying during mild and moderate blood loss but become overwhelmed once a critical reduction in circulating volume is reached. The major stratified decreases in EDP across all hemorrhage severities, bolstering the idea that ventricular filling pressures were greatly impacted, despite declining systolic performance, likely reflecting coordinated cardiovascular compensation. This negative impact on cardiac performance dynamics was mirrored by systemic reductions in renal and carotid perfusion, which were also significantly impacted by increasing hemorrhage levels. The parallel decline in renal and carotid flow underscores the global nature of the hypovolemic insult at the end of hemorrhage.

At T60, both hemorrhage severity and occlusion type had significant effects on cardiac mechanics, revealing elevated physiological stress. Increases in Ea, ESV and ESP were paired with by reductions in EDV and SW. This implies occlusion both decreased preload and increased afterload conditions, challenging ventricular emptying. Additionally, parameters that reflected the limits of systolic compensation, namely SV, SW and EJF, declined most prominently under severe hemorrhage or occlusion, highlighting the constrained capacity of the heart to adapt under the combined impact of these stressors. These ultimately illustrate that while occlusion may stabilize proximal pressures, it does so at the cost of increased ventricular workload and altered loading conditions.

In contrast to the perfusion decline relative to increasing hemorrhage levels observed at T30, renal and carotid blood flow behavior was divergent at T60, reflecting regional perfusion redistribution as a function of occlusion. Renal arterial flow was inhibited by increasing occlusion. For instance, full occlusion produced severe and consistent reductions in renal flow through all hemorrhage levels. This consistent trend demonstrates that occlusion, independent of hypovolemic state, was detrimental to renal flow, an effect that was expected considering the flow mechanics induced by full occlusion.

Conversely, carotid flow was promoted with increasing occlusion, consistent with elevated cerebral circulation^16^. With increased occlusive stress, however, augmented carotid perfusion effect was stunted by an increase in hemorrhage severity, demonstrating that cerebral flow preservation capacity is constrained by available blood volume. The interaction between hemorrhage severity and occlusion strategy was statistically significant for carotid flow, highlighting the dynamic balance between perfusion redistribution and hemorrhagic blood volume depletion during resuscitative occlusion.

These results emphasize the typical inherent physiological exchange from REBOA implementation^15,21–23^. The preservation of proximal and cerebral perfusion occurs in the system at the expense of distal organ blood flow, like in the case of the kidneys, paired with increases in cardiac stress. This is demonstrated by the P-V data which provides mechanistic insight into this trade-off where occlusion elevates afterload while impairing ventricular emptying, while concurrently reducing stroke work. These physiological costs likely contribute to the morbidity associated with prolonged or complete occlusion and help explain the variability observed in clinical outcomes.

Additionally, the study’s findings support the rationale for partial REBOA strategies. For instance, while f-REBOA demonstrates increases in both proximal pressure and carotid perfusion, it simultaneously induces renal hypoperfusion and imposes substantial cardiac load, promoting cardiac injury. p-REBOA, conversely, maintains a pressure gradient across the REBOA balloon while in the system, affording a way to mitigate risk of supraphysiological afterload elevation found in f-REBOA and reducing the risk of distal ischemia in regions like the kidney, all while supporting proximal cerebral perfusion. These responses were similarly observed across all hemorrhage levels, further promoting this benefit. This trend matches what was observed in previous studies performed by our group^15^.

While the present analysis focused on these left ventricular P-V relationships and renal and carotid perfusion to validate the model against established physiological responses detailed in the literature, the extent of the collected dataset supports a wide range of further investigation. This dataset enables future analyses that can functionally integrate additional study into aortic and venous flow and pressure across the remaining vascular beds in conjunction with the incremental systemic blood gas measurements, biomarkers of ischemia–reperfusion injury, urine output, and post-mortem histology to further investigate the systemic and cellular consequences of varying occlusion strategies in conjunction with varying levels of hemorrhage. Additionally, the nature of the randomized design of this model permits further analysis into the interaction between hemorrhage severity, occlusion modality, and post-injury fluid and drug resuscitation strategy to pave the initial steps for a translational framework to optimize clinical translatability beyond REBOA employ.

Several limitations to this study should be acknowledged. First, the data gathered corresponds to a controlled hemorrhage large-animal model under anesthesia, meaning the rate of blood loss was predefined and constant, limiting translatability to uncontrolled hemorrhage settings. However, our team has experience with implementing an uncontrolled liver laceration hemorrhage model within this translational porcine model as presented by Cambronero/Sanin et al, where p-REBOA also outperformed other conventional REBOA strategies. Secondly, we recognize the limitation of a sternotomy within our pig model which may have reduced our starting systemic pressures, along with the use of isoflurane. Future studies could incorporate less invasive pressure-volume catheters via Swan Ganz or alternative methods to capture cardiac performance such as doppler ultrasound or 4D MRI. Furthermore, while high-resolution pressure-volume analysis gave us detailed insight into left ventricular mechanics, this study focused primarily on load-dependent parameters and no inferior vena caval occlusion protocol to gather load-independent data.

## Conclusion

In this study we developed and exploited a novel translational porcine model with automated control to assess the physiologic responses to varying degrees of hemorrhage and aortic occlusion. Continuous hemodynamic data was captured from four pressure probes and six flow probes including measurements of carotid and renal arterial flow, left ventricular cardiac performance parameters and intravenous drug and fluid administration. We demonstrated a graded decline in cardiac performance and systemic perfusion with increasing hemorrhage severity and noted periods of compensatory vs. decompensatory responses. With respect to aortic occlusion, 30 minutes of f-REBOA resulted in significant ischemia-reperfusion injury where renal perfusion was profoundly suppressed to 40% of baseline renal flow. On the other hand, p-REBOA yielded superior renal perfusion, while maintaining cardiac function and carotid perfusion. p-REBOA also required less fluid and vasopressor use during the critical care period. Together, this unique dataset forms the basis for interpreting the integrated physiological response to hemorrhage, REBOA interventions, and subsequent resuscitation, providing new opportunities to assess acute cardiovascular hemodynamics for trauma and acute critical care.

## Acknowledgements

We’d like to acknowledge Hebah Soudan and Magan Lane for their assistance with some of the animal experiments in this study. We’d also like to acknowledge Mr. Jerry Louis Shelton at Texas A&M University for his assistance in generating the pig illustration in Figure 1A.

## References

1. Schiller AM, Howard JT, Convertino VA. The physiology of blood loss and shock: New insights from a human laboratory model of hemorrhage. Exp Biol Med. 2017;242(8):874–883. doi:10.1177/1535370217694099

2. Johnson AB, Burns B. Hemorrhage. In: StatPearls. StatPearls Publishing; 2024. Accessed September 16, 2024. http://www.ncbi.nlm.nih.gov/books/NBK542273/

3. Latif RK, Clifford SP, Baker JA, et al. Traumatic hemorrhage and chain of survival. Scand J Trauma Resusc Emerg Med. 2023;31:25. doi:10.1186/s13049-023-01088-8

4. Eastridge BJ, Holcomb JB, Shackelford S. Outcomes of traumatic hemorrhagic shock and the epidemiology of preventable death from injury. Transfusion (Paris). 2019;59(S2):1423–1428. doi:10.1111/trf.15161

5. Kauvar DS, Lefering R, Wade CE. Impact of hemorrhage on trauma outcome: an overview of epidemiology, clinical presentations, and therapeutic considerations. J Trauma. 2006;60(6 Suppl):S3–11. doi:10.1097/01.ta.0000199961.02677.19

6. Brenner M. The Role of Resuscitative Endovascular Balloon Occlusion of the Aorta. Surg Clin North Am. 2024;104(2):311–323. doi:10.1016/j.suc.2024.01.003

7. King DR. Initial Care of the Severely Injured Patient. N Engl J Med. 2019;380(8):763–770. doi:10.1056/NEJMra1609326

8. Thrailkill MA, Gladin KH, Thorpe CR, others. Resuscitative Endovascular Balloon Occlusion of the Aorta (REBOA): update and insights into current practices and future directions for research and implementation. Scand J Trauma Resusc Emerg Med. 2021;29(1):8. doi:10.1186/s13049-020-00807-9

9. Ordonez CA, Parra MW, Caicedo Y, others. REBOA as a New Damage Control Component in Hemodynamically Unstable Noncompressible Torso Hemorrhage Patients. Colomb Med Cali. 2020;51(4):e4064506. doi:10.25100/cm.v51i4.4422.4506

10. Joseph B, Zeeshan M, Sakran JV, et al. Nationwide Analysis of Resuscitative Endovascular Balloon Occlusion of the Aorta in Civilian Trauma. JAMA Surg. 2019;154(6):500–508. doi:10.1001/jamasurg.2019.0096

11. Burkhoff D, Mirsky I, Suga H. Assessment of systolic and diastolic ventricular properties via pressure-volume analysis: a guide for clinical, translational, and basic researchers. Am J Physiol Heart Circ Physiol. 2005;289(2):H501–512. doi:10.1152/ajpheart.00138.2005

12. Mirsky I, Tajimi T, Peterson KL. The development of the entire end-systolic pressure-volume and ejection fraction-afterload relations: a new concept of systolic myocardial stiffness. Circulation. 1987;76(2):343–356. doi:10.1161/01.cir.76.2.343

13. van der Velde ET, Burkhoff D, Steendijk P, Karsdon J, Sagawa K, Baan J. Nonlinearity and load sensitivity of end-systolic pressure-volume relation of canine left ventricle in vivo. Circulation. 1991;83(1):315–327. doi:10.1161/01.cir.83.1.315

14. Claessens TE, Georgakopoulos D, Afanasyeva M, et al. Nonlinear isochrones in murine left ventricular pressure-volume loops: how well does the time-varying elastance concept hold? Am J Physiol-Heart Circ Physiol. 2006;290(4):H1474–H1483. doi:10.1152/ajpheart.00663.2005

15. Fu M, Ac R, E CP, et al. Investigating the variability in pressure-volume relationships during hemorrhage and aortic occlusion. PubMed. Published online 2023. Accessed July 24, 2025. https://pubmed.ncbi.nlm.nih.gov/37680564/

16. Matsumura Y, Higashi A, Izawa Y, others. Distal pressure monitoring and titration with percent balloon volume: feasible management of partial resuscitative endovascular balloon occlusion of the aorta (P-REBOA). Eur J Trauma Emerg Surg. Published online 2019. doi:10.1007/s00068-019-01257-4

17. Sanin GD, Cambronero GE, Wood EC, et al. MAN VERSUS MACHINE: PROVIDER DIRECTED VERSUS PRECISION… : Shock. Accessed July 24, 2025. https://journals.lww.com/shockjournal/fulltext/2024/05000/man_versus_machineprovider_directed_versus.14.aspx

18. Stonko DP, Edwards J, Abdou H, et al. A technical and data analytic approach to pressure-volume loops over numerous cardiac cycles. JVS-Vasc Sci. 2022;3:73–84. doi:10.1016/j.jvssci.2021.12.003

19. Reva VA, Matsumura Y, Hörer T, et al. Resuscitative endovascular balloon occlusion of the aorta: what is the optimum occlusion time in an ovine model of hemorrhagic shock? Eur J Trauma Emerg Surg. 2016;44(4):511–518. doi:10.1007/s00068-016-0732-z

20. Kuckelman JP, Barron M, Moe D, others. Extending the golden hour for Zone 1 resuscitative endovascular balloon occlusion of the aorta: Improved survival and reperfusion injury with intermittent versus continuous resuscitative endovascular balloon occlusion of the aorta of the aorta in a porcine severe truncal hemorrhage model. J Trauma Acute Care Surg. 2018;85(2):318–326. doi:10.1097/TA.0000000000001964

21. Russo RM, White JM, Baer DG. Partial Resuscitative Endovascular Balloon Occlusion of the Aorta: A Systematic Review of the Preclinical and Clinical Literature. J Surg Res. 2021;262:101–114. doi:10.1016/j.jss.2020.12.054

22. Johnson MA, Davidson AJ, Russo RM, others. Small changes, big effects: The hemodynamics of partial and complete aortic occlusion to inform next generation resuscitation techniques and technologies. J Trauma Acute Care Surg. 2017;82(6):1106–1111. doi:10.1097/TA.0000000000001446

23. Davidson AJ, Russo RM, DuBose JJ, others. Potential benefit of early operative utilization of low profile, partial resuscitative endovascular balloon occlusion of the aorta (P-REBOA) in major traumatic hemorrhage. Trauma Surg Acute Care Open. 2016;1(1):e000028. doi:10.1136/tsaco-2016-000028

